# Using AlphaFold for Rapid and Accurate Fixed Backbone Protein Design

**DOI:** 10.1101/2021.08.24.457549

**Authors:** Lewis Moffat, Joe G. Greener, David T. Jones

## Abstract

The prediction of protein structure and the design of novel protein sequences and structures have long been intertwined. The recently released AlphaFold has heralded a new generation of accurate protein structure prediction, but the extent to which this affects protein design stands yet unexplored. Here we develop a rapid and effective approach for fixed backbone computational protein design, leveraging the predictive power of AlphaFold. For several designs we demonstrate that not only are the AlphaFold predicted structures in agreement with the desired backbones, but they are also supported by the structure predictions of other supervised methods as well as *ab initio* folding. These results suggest that AlphaFold, and methods like it, are able to facilitate the development of a new range of novel and accurate protein design methodologies.

## 1 Introduction

The computational design of proteins provides a route to create new protein structures and functions (Kuhlman and Bradley, 2019), with applications such as therapeutics (Chevalier et al., 2017), nanomaterials (King et al., 2012), and enzymology (Studer et al., 2018) amongst others. *De novo* protein design has historically been carried out using energy functions (Alford et al., 2017; Huang et al., 2020) and extensive sampling, which has been successful but has high resource requirements and a high rate of failure. Recently deep learning techniques have impacted both protein structure prediction and protein design (Pearce and Zhang, 2021; Wittmann et al., 2021). Design tasks tackled with deep learning include fixed backbone design (O’Connell et al., 2018; Ingraham et al., 2019; Qi and Zhang, 2020; Norn et al., 2021), antibody design (Wang et al., 2018; Saka et al., 2021; Shin et al., 2021), *de novo* design (Anishchenko et al., 2020;Moffat and Jones, 2021), and the prediction of whether a sequence has a stable structure from sequence alone (Singer et al., 2021). A variety of neural network architectures have been used including variational autoencoders (Greener et al., 2018; Hawkins-Hooker et al., 2021), deep exploration networks (Linder et al., 2020), graph neural networks (Strokach et al., 2020), recurrent neural networks (Alley et al., 2019) and autoregressive models (Shin et al., 2021;Trinquier et al., 2021). Ultimately the hope is that faster and more accurate protein design with deep learning will lead to the design of functional proteins (Tischer et al., 2020; Caceres-Delpiano et al., 2020).

Historically, improvements in protein structure prediction have led to improvements in protein design. The trRosetta structure prediction method has been used to design a variety of proteins by modifying random starting sequences to give sharp predicted residue-residue distance maps (Anishchenko et al., 2020). A related approach used the gradients of the trRosetta network to optimise sequences for a given backbone (Norn et al., 2021). The AlphaFold method from DeepMind performed exceptionally well at the Critical Assessment of protein Structure Prediction (CASP) 14 assessment, effectively solving the structure prediction problem for monomeric domains with evolutionary information available (Jumper et al., 2021). Recently AlphaFold has been made available open source and the community has already modified it to predict protein complexes, a task for which it was not designed (Mirdita et al., 2021). It is clear that AlphaFold has captured a complex and intricate understanding of protein structure prediction. What is not yet immediately clear is the extent to which this can be leveraged in the opposite direction for protein design. Protein structure prediction methods tend to perform well on designed proteins in the absence of coevolutionary information (Xu et al., 2021) and it is reasonable to expect that within AlphaFold is the potential for rapid and accurate protein design.

Here we explicitly demonstrate that potential by developing a technique for computational protein design using AlphaFold. More specifically, we develop an approach for fixed backbone design that leverages multiple features of AlphaFold to efficiently design sequences that are strongly predicted to fold to a specified backbone. This is done by taking a starting sequence, which is biased towards a solution by sampling from a generative sequence model of *de novo* proteins, and iteratively mutating it to find a final sequence that has a predicted structure that fits the desired backbone with high confidence. We find the AlphaFold predicted structures for the designed sequences are supported by predictions from other supervised protein structure predictors as well as *ab initio* folding with Rosetta (Leaver-Fay et al., 2011).

## 2 Methods

In this section we describe how we adapt AlphaFold to perform protein design using end-2-end structure prediction. We first discuss how we generate a starting sequence, with the aim of starting the design process as close as possible to a final solution. We then go on to describe the iterative design process itself followed by the computational evaluation of the designed sequences.

### 2.1 Generating a starting sequence design

The first step in the design process is generating a starting sequence. Instead of beginning with a sequence of randomly chosen amino acids, or a simple polyalanine chain, we leverage a method currently under development, consisting of a generative sequence model (Moffat and Jones, 2021), to kick-start the design process. This generative model consists of an autoregressive transformer model (Radford et al., 2019; Vaswani et al., 2017), via causal masking, trained on several hundred thousand computationally de novo designed sequences (Moffat and Jones, 2021). We sample 1,000 sequences of *L* = 100, where L is length of a sequence in residues, from the generative model. These samples are run through the inference pipeline of AlphaFold and we keep the best relaxed structure as chosen by the highest pLDDT (the model’s predicted score on the lDDT-C*α* metric, more details in Jumper et al. (2021)) in each case. This provides us with a small sequence and predicted structure database, consisting of 1,000 sequence & predicted structure pairs, which we then use to construct the starting sequence.

For the fixed backbone design task we have some backbone we wish to design a sequence for. In this work we explore designs for several backbones, originating from previously designed and validated *de novo* designed proteins. Assuming we have some specific backbone we wish to design for, we take the coordinates file of the backbone and compare it to every structure in the previously constructed database using TMAlign (Zhang and Skolnick, 2005).

We take the database structure with the highest predicted TM-score (Zhang and Skolnick, 2005) relative to the backbone candidate and use the structural alignment to choose which residues from the database structure will be used to construct the starting sequence. Residues in the database sequence that are aligned to the backbone structure are kept to be part of the starting sequence. Residues that are not aligned are removed. Any gaps in the database sequence are replaced with alanine. This process is visualized in Figure 1.

**Figure 1:**
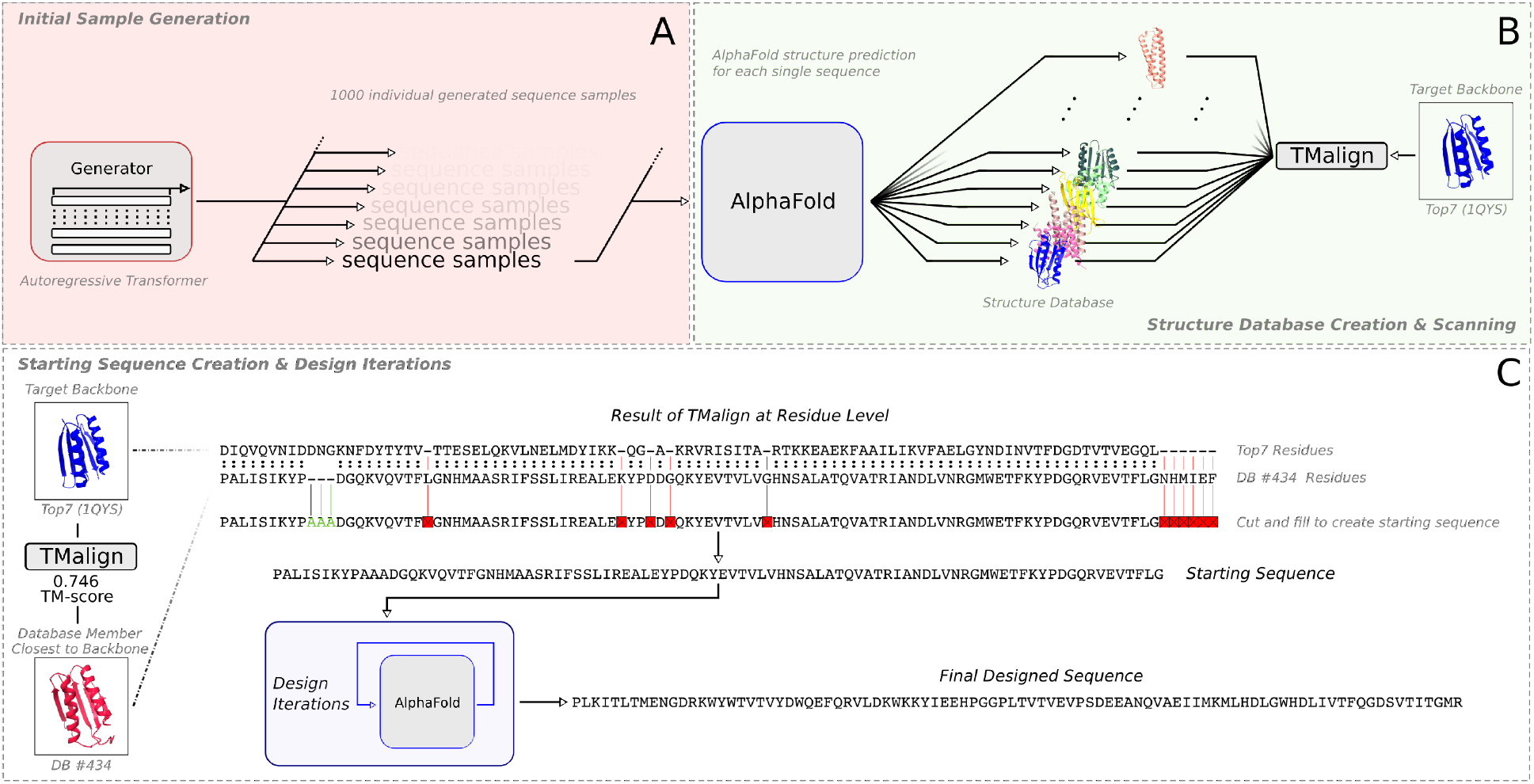
An overview of the fixed backbone design pipeline. (**A**) 1,000 individual sequence samples are drawn from the generative model. (**B**) Each individual sequence has its structure predicted by AlphaFold, with the best ranked structure by pLDDT score being kept as the final predicted structure in each case. Together these make a database that is scanned against with TMAlign and the desired backbone. (**C**) The sequence and predicted structure with the highest TM-score to the backbone are taken and used to create a starting sequence.

This approach produces a starting sequence that is the same length as the backbone by utilizing residue fragments from sequence samples that have been produced by the generative sequence model. This has the advantage of biasing the starting sequence towards a residue composition that is more likely to lead to the correct predicted backbone structure than if the residues were simply randomly chosen.

### 2.2 Iterative end-2-end design

We design sequences with AlphaFold by taking the starting sequence and iteratively improving it until it produces a predicted structure matching the desired backbone. At a high level, this is done by performing a greedy semi-random walk through sequence space, progressively mutating the starting sequence. The mutations are chosen so as to minimize the cross-entropy between the pseudo-C*β* distogram predictions and the pseudo-C*β* distances of the backbone. The term pseudo-C*β* is used as C*α* coordinates are used for glycine residues and C*β* for all others.

More concretely, beginning with the starting sequence, we predict its structure with AlphaFold and the distogram based loss. This loss is the same implementation of distogram loss as used in training AlphaFold (Jumper et al., 2021). In order to benefit from ensembling, predictions are made with each set of parameters (each set dubbed a model in the AlphaFold documentation) *M* used in CASP14. We use parameter set and model interchangeably. This produces *M* = 5 sets of results as five different parameters sets were used. The scalar loss from each individual parameter set *m* is averaged together for a final distogram loss 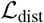,

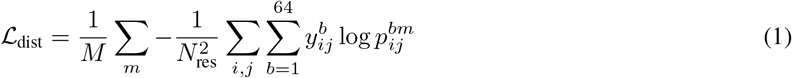

where *i, j* are the row and column of the distogram prediction *p*, and *y*, which is the ground-truth distance binned into one of 64 bins *b* at each *i, j* position. The pLDDT is also calculated during inference and then averaged across parameter sets but not across sequence length. These averaged pLDDT predictions are used to calculate weights *W* = {*W*_1_,…, *W_L_*} at each position, *i*. The normalized weights are then treated as probabilities from a discrete distribution over sequence positions from which we sample a position.

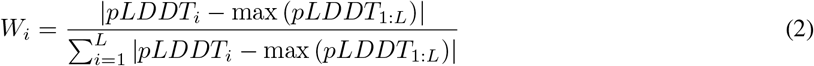

Clearly, this weighting approach favours positions with low pLDDT scores and disfavours positions with high scores. By biasing the position selection in this way, the method attempts to improve the sequence prediction’s fit to the backbone where AlphaFold is least confident with the prediction. Amino acids in the sequence that lead to low confidence predictions are less likely to lead to a stable structures. Thus, assuming that the desired target backbone is itself stable, a mutation in a low confidence region is more likely to result in an improvement to the loss.

After the position is sampled, the amino acid at the sampled position is randomly mutated into another, uniformly sampled from all other amino acids with the exception of cysteine. AlphaFold predictions are generated for the mutated sequence and if and only if the mutated sequence improves the distogram loss then the mutant is kept, replacing the starting sequence.

Overall, this constitutes one design iteration. For each target backbone design explored in this work, we perform 20,000 design iterations. This is equivalent to testing 20,000 individual mutants. In all cases explored this was sufficient for the distogram score to have converged. The current sequence after performing the full steps is then taken forward as the final designed sequence for computational evaluation.

### 2.3 Fast AlphaFold inference

In order to use AlphaFold to design protein sequences we need to be able to perform inference a large number of times and so consequently at a reasonable speed. We configure the publicly available AlphaFold implementation to perform inference on a single sequence without evolutionary information in under a second on consumer-grade RTX GPUs (e.g. RTX 30 and 20 series) using one parameter set. Due to availability, a variety of RTX graphics cards were used for generating designs, however, this was most commonly an RTX 2080 Ti, which we use for benchmarking. Inference using one model takes 0.9s, and one complete design iteration, which uses all 5 parameter sets, takes roughly 5 s, for sequences around 100 residues long.

During design and evaluation we use slightly different configurations but in both cases the same configuration is used across all parameter sets. We highlight this point as the default configurations differ between the 5 parameter sets. During design we disable template calculations and reduce the maximum sequences used in the MSA portion of the Evoformer trunk down to 1. All recycling is also disabled. All heads that are not used as a part of the design process are disabled. When we use AlphaFold for evaluation we keep this configuration the same but increase the number of recycles performed to 10. We also only perform relaxation (Eastman et al., 2017; Jumper et al., 2021) during evaluation consistent with the default AlphaFold pipeline.

### 2.4 Computational evaluation of designed sequences

The first type of computational approach we use to evaluate and validate a designed sequence is to assess the fit of its predicted structure by AlphaFold to the target backbone. This primarily constitutes running AlphaFold as discussed previously and taking the relaxed structure with the highest pLDDT score as the predicted structure. We also compare against the predicted structure from trRosetta, a deep supervised structure predictor (Yang et al., 2020), that has been used recently for structure prediction in design tasks (Norn et al., 2021).

The second approach we take to evaluating the sequence is a more traditional protein design approach. More specifically, we perform *ab initio* structure prediction with Rosetta (Leaver-Fay et al., 2011). First, fragments are generated for the designed sequence using the Robetta (Kim et al., 2004) server without homologues. On a local HPC cluster, 15,000 decoys are then generated using the Rosetta Abinitio Relax protocol (Raman et al., 2009). We take the best decoy by Rosetta all atom score (Alford et al., 2017) as the predicted structure. We also perform homology searches for the designed sequences against UniRef100 (June-2021) using MMseqs2 (Steinegger and Söding, 2017) with a sensitivity parameter of 7.0 and 1 iteration. For visualization of protein structures we use ChimeraX (Pettersen et al., 2021).

## 3 Results

### 3.1 Rapid Fixed Backbone Design

As the first example of this method, we generate a design for the backbone of Top7 (Kuhlman et al., 2003). Beginning with construction of the starting sequence, the Top7 backbone has a TM-Score of 0.746 with the closest member of the database. This similarity to the database member can be seen in Figure 1. After generating the starting sequence and subjecting it to design, the final designed sequence shares 27%sequence identity with the original Top7 sequence (Figure 2). The AlphaFold predicted structure of the design has a C*α*-RMSD of 0.736 Å from the Top7 backbone. AlphaFold also produces a high confidence prediction with a pLDDT score of 91.

**Figure 2:**
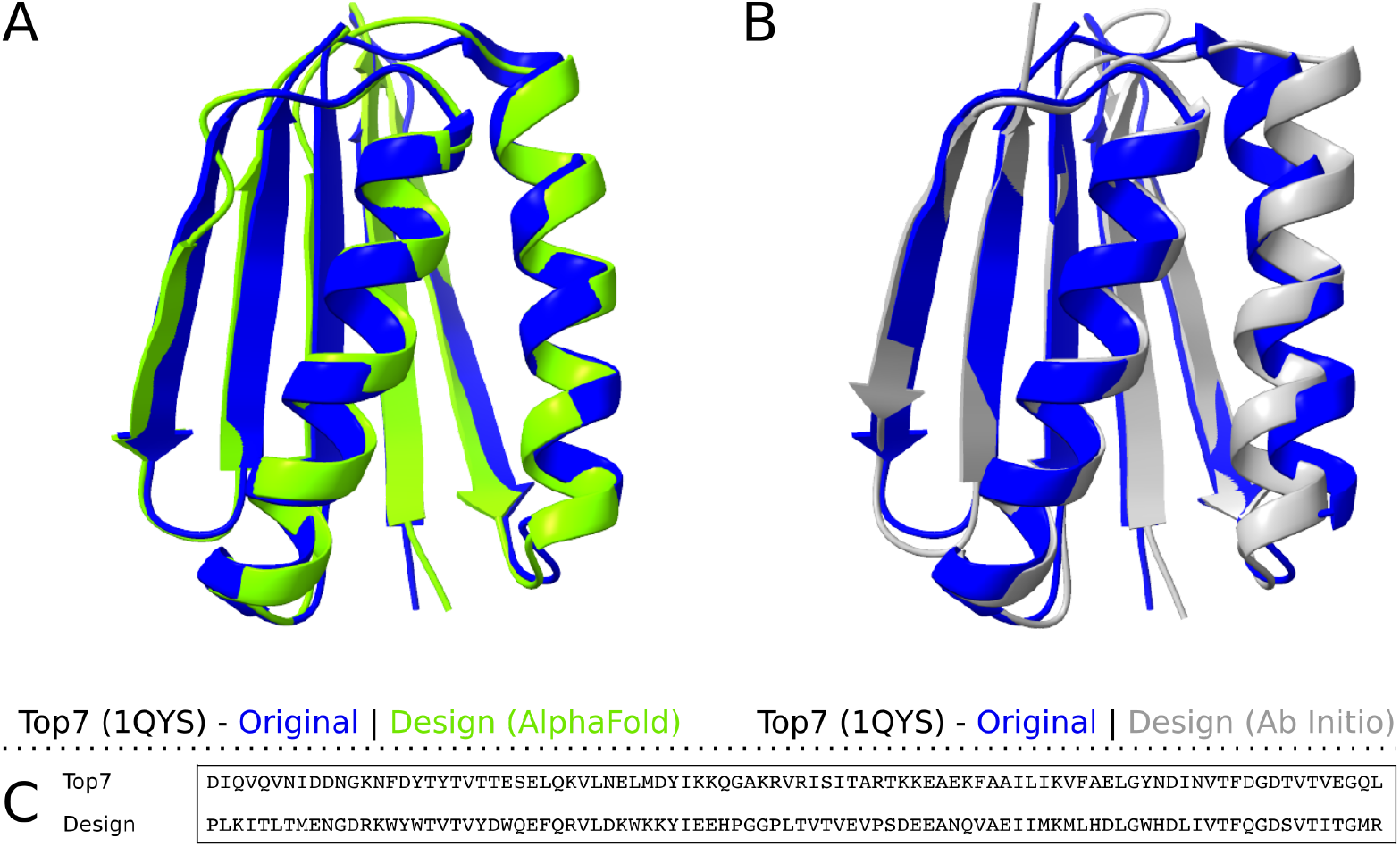
The superimposed structures of Top7 (PDB ID 1QYS) and the predicted structures of the new design, being the sequence newly designed to fold to Top7’s backbone. (**A**) The structure prediction of the Design as generated by AlphaFold. The prediction has a C*α*-RMSD of 0.736 Å relative to Top7. (**B**) The structure prediction of the Design generated by Rosetta *ab initio* folding simulations. The prediction shown is the best decoy by the Rosetta all atom score, being –297. It has a C*α*-RMSD of 0.736 Å relative to Top7. (**C**) The sequences of the Design and Top7. The Design has a sequence identity of 27% with Top7.

The AlphaFold prediction is also supported by the trRosetta prediction. The trRosetta best predicted structure is also very close to the desired backbone, with a C*α*-RMSD of 2.637 Å. When trRosetta performed the prediction it found no homologues relative to Uniclust30 (Mirdita et al., 2017), predicting from only the single sequence, and it returned a high confidence prediction with an estimated TM-score of 0.679. Both supervised methods producing highly similar and highly confident results that fit well to the desired backbone suggests that the designed sequence is likely to fold to the desired backbone.

This is further supported by the results from having performed *ab initio* folding simulations with Rosetta. The best decoy by the Rosetta all atom score (−297), from the 15,000 generated, has a C*α*-RMSD of 1.279 Å from the backbone (Figure 2). This strongly suggests that the designed sequence will not only fold, but importantly, fold to the backbone. Running MMseqs2 search against UniRef100 (Suzek et al., 2015) yielded 2 hits. The first is, unsurprisingly, to the original Top7 (PDB ID 1QYS) sequence. The second is to a mutant of Top7, 2N41 by PDB ID, that has an 88% sequence identity to Top7. This mutant also has a sequence identity of 27% to the designed sequence. This suggests that the designed sequence represents a novel solution.

### 3.2 Designing with Unseen Backbones

The results for designing a sequence folding to the backbone of Top7 are very encouraging. That being said, the design provides a limited estimation of the potential of this approach as it is likely that the original Top7 was present in the training set of AlphaFold. It is difficult to quantify what, if any, effect this has on the design process, but it is possible that this provides some bias towards designing a sequence similar to the original Top7 sequence, which is itself more likely to fold correctly. Practically, the clearest approach to avoiding this issue is to create designs for recently released backbones not present in the AlphaFold training set. This ensures that AlphaFold does not have a bias towards the backbone’s original sequence due to not having seen either backbone or sequence during training as an example. To that end, we design sequences that are confidently predicted to fold to the desired backbones, by AlphaFold and *ab initio* folding, for Peak6 (PDB ID 6MRS), Foldit1 (6MRR) and Ferredog-Diesel (6NUK), all released after the cutoff date (30 April 2018) for the AlphaFold training set.

#### 3.2.1 Peak6

The first protein backbone we design to is Peak6 (PDB ID 6MRS). As was done in the last example, we take Peak6 and use TMAlign to align and score it against each of the 1,000 members of the database, where each member is an AlphaFold predicted structure of a sequence sampled from a generative model. The closest database member to the Peak6 backbone has a TM-score of 0.596 and it is used to construct a starting sequence as illustrated in Figure 1. AlphaFold produces a very confident prediction for the final designed sequence with a pLDDT score of 95, and the prediction itself has a C*α*-RMSD of 0.537 Å from the target backbone. The designs for Peak6, and the other designs discussed in this section can be seen in Figure 3 along with the original backbone, the original backbone’s amino acid sequence, and the amino acid sequence of the design. The results of *ab initio* folding also result in a good fit to the backbone, with a Rosetta all atom score of −244 and a C*α*-RMSD Å of 2.301. The designed sequence has a sequence identity of 27% to the original Peak6 sequence which is very encouraging given that AlphaFold has not seen Peak6 as part of training.

**Figure 3:**
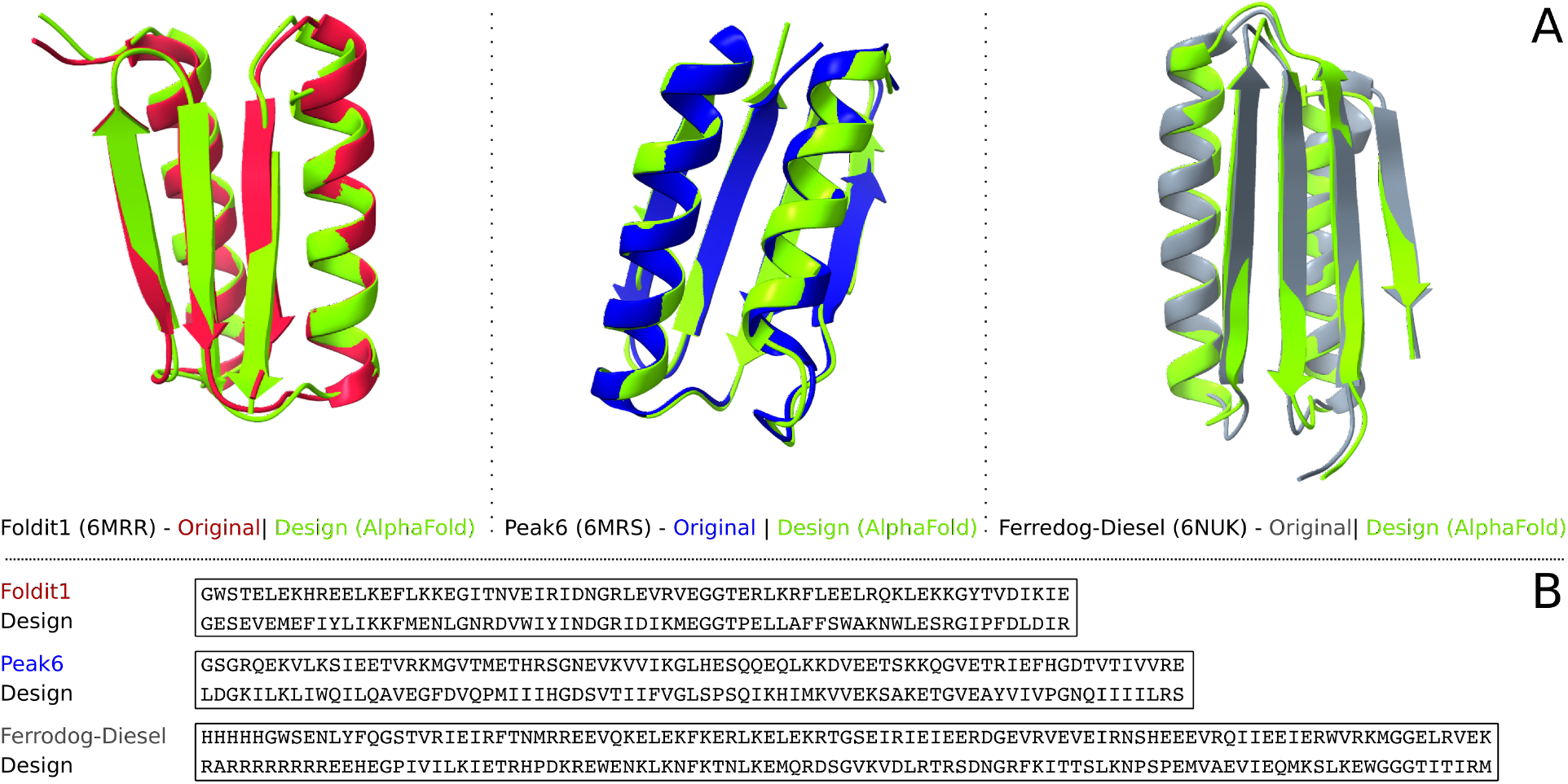
(**A**) AlphaFold predicted structures along with the original desired backbone for three design cases. For Ferredog-diesel we do not include the first 16 residues the long N-terminal tail for clarity. (**B**) Sequences of the original backbones in **A** along with the sequences designed in this work to fit to the backbones. Residues in the original backbone sequence, at the N-terminus and C-terminus, that do not have coordinates reported in the original PDB file are not displayed. For Ferredog-diesel, the first 16 residues are not included in any sequence identity calculation. The sequence identity between the designed sequence and the original sequence for Foldit1, Peak6, and Ferrodog-Diesel, is 31%, 27%, and 22% respectively.

trRosetta produces a confident prediction with an estimated TM-score of 0.618 however the prediction diverges from the backbone, having a C*α*-RMSD of 11.444 Å. Upon inspection this was due to trRosetta having constructed a large multiple sequence alignment (MSA) with 7725 weak hits, generated by HHblits(Remmert et al., 2012) using Uniclust30 (08-2018), which biased the prediction. Corroborating this, running MMseqs2 search against UniRef100 generated no hits. Using HHblits to search against a more recent Uniclust30 (06-2021) version with default parameters results in 94 weak hits, with the lowest having an E-value of 0.9, being A0A2N5PTC8, a small uncharacterized protein. The second lowest is the original Peak6 sequence with an E-value of 0.95. As further confirmation of a good design we predict its structure with DMPfold2 (Kandathil et al., 2021) without evolutionary information. As is expected, it returns a prediction that is a good fit to the backbone with a C*α*-RMSD of 2.055 Å.

For the Peak6 designed protein, we plot the trajectory of the distogram loss that is being optimized during design, alongside the pLDDT and the sequence identity to the original backbone sequence. This can be seen in Figure 4. All three metrics shown quickly improve during the early stages of design then continue on to plateau. Although the sequence identity does not appear to significantly improve after 5,000 iterations, the loss and pLDDT continue to slowly improve. This suggests that, during design, the designed sequence rapidly approaches the correct backbone structure early on, and that the majority of proceeding design steps only make small improvements to the fit.

**Figure 4:**
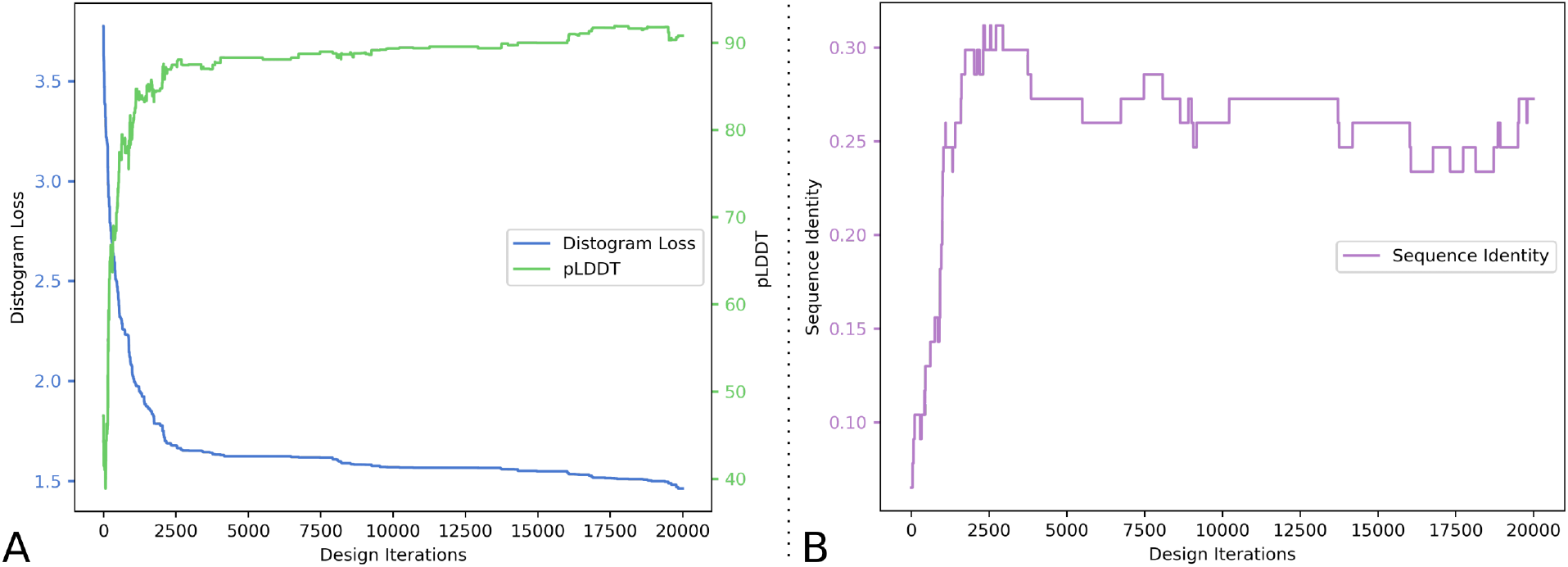
The change in several metrics over the course of designing the sequence that is predicted to fold to the Peak6 (6MRS) backbone. (**A**) A plot showing the change in Distogram Loss 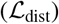 on the left-axis, and change in pLDDT on the right-axis, with design iterations. (**B**) A plot showing the change in sequence identity between the designed sequence at each iteration and the original Peak6 sequence, with design iterations.

#### 3.2.2 Foldit1

The next sequence we design is for the Foldit1 backbone (PDB ID 6MRR). The closest database member has a TM-score of 0.517 with the backbone, and after design AlphaFold produces a confident prediction that has a pLDDT score of 88 and a C*α*-RMSD of 0.888 Å. The designed sequence also shares an encouraging 31% sequence identity with the original Foldit1 sequence. Similarly to the previous design of Peak6, trRosetta found and included 691 weak hits found by HHblits leading to a C*α*-RMSD of 13.184 Å from the backbone. By comparison, DMPfold2 produces a prediction, without using evolutionary information, with a C*α*-RMSD of 3.815 Å. Running HHblits and MMseqs2 search in the same fashion as previously results in MMseqs2 finding no hits and HHblits only returning 95 hits. Of the 95 hits, 94 have an E-value of ≥ 0.16 and mostly constitute weak hits to uncharacterized proteins and protein fragments. The one strong hit has an E-value of 5.4e-4 and is the original Foldit1 sequence. This is positive evidence that the designed sequence is likely to fold to the desired backbone. Even further evidence is provided in that the best *ab initio* decoy, with a Rosetta all atom score of −205, has a C*α*-RMSD 1.96 Å to the backbone.

#### 3.2.3 Ferredog-Diesel

The sequence designed for Ferredog-Diesel (PDB ID 6NUK), like the previous designs, is also well predicted to fold to the desired backbone. During the design we include the long N-terminal tail but for the values reported here we ignore it by not including the first 16 residues in the given calculations. The database member closest to the Ferredog-Diesel backbone has a TM-score of 0.517 and the final designed sequence, as expected, has an AlphaFold predicted structure with a high confidence and close fit the the backbone, the pLDDT score being 84 and the C*α*-RMSD being 0.990 Å.

Both trRosetta and DMPfold2 also provide predictions, using only the single sequence, with good fits to the desired backbone, the C*α*-RMSD being 1.723 Å and 2.260 Å respectively. To maintain consistency with the previous two examples we perform a sequence search with HHblits in addition to MMseqs2 search. While MMseqs2 finds no hits, HHblits finds hits resulting in a similar outcome to the Foldit1 designed sequence. It finds 118 hits, all with E-values of ≥ 0.0.57 with the exception of 1 hit, the original Ferredog-Diesel sequence, which has an E-value 0.039. The sequence identity between the original Ferredog-Diesel sequence and the designed sequence is 22%. The sequence identity and sequence search results are also supportive of the designed sequence folding to the desired backbone. The results of *ab initio* folding are also supportive of the design, with a C*α*-RMSD of 2.467 Å and a Rosetta all atom score of −283.

## 4 Discussion

Here we present a conceptually simple approach to fixed backbone protein design that utilizes the flexibility and high accuracy of end-2-end structure prediction provided by AlphaFold (Jumper et al., 2021). From a protein structure prediction perspective, end-2-end methods are clearly very attractive, even discounting performance. They consolidate the process of going from sequence to structure to one forward pass through a neural network. Given structure prediction’s role in protein design this opens up many potentially fruitful avenues for their use in novel protein design approaches.

A very useful aspect of the AlphaFold model is its use of multiple prediction heads. In this work we utilize both the pLDDT scores and the distogram predictions as part of a greedy optimization approach that can successfully design sequences that are predicted to fold to the desired backbone *in silico*. For the examples explored here, it is very encouraging that the solutions converged upon by AlphaFold were supported by other predictors, and especially by *ab initio* folding. One of the clear benefits to this approach is its stark simplicity. Not only does this open up different ways to extend the method, it facilitates easy investigation during the design process itself. At each step, the current sequence is available along with an all-atom prediction of its structure, a probabilistic prediction of the backbone structure, a probabilistic prediction of the residues that could occupy each sequence position, a per residue confidence score in the pLDDT, and more. At any point in the design process, these predictions are available to be investigated and probed by the human designer if they so choose.

Another especially interesting aspect of the approach presented here is the absence of an explicit energy function in the design process, excluding the final relaxation (Eastman et al., 2017) performed as a part of evaluation, as is a staple of the most advanced current protein design approaches (Marcos et al., 2018; Vorobieva et al., 2021). Given that this approach produces designs that are unique on a sequence level, matching a backbone it has not been trained on, and that it assigns a high confidence, it would speculatively suggest that AlphaFold has learned some generalization of protein structures that relates to an energy function. As AlphaFold and similar methods continue to be used, the relationship between the structure predictor and current energy function based folding approaches may prove a promising avenue for future research.

Although there is a large amount of stochasticity in the selection of mutations it is nonetheless interesting that a greedy approach is effective. Going forward, there is still a risk of designed sequences being trapped in local minima and so looking to develop alternative non-greedy optimization approaches may be a valuable first step in looking to improve the current method. One immediate option would be to explore an optimization approach more likely to converge to a global minimum like simulated annealing or incorporating mutations at multiple locations in a given step. Another alternative approach is gradient optimization of the distogram loss. A gradient descent approach is likely to also be susceptible to becoming trapped in local minima but it would hopefully reduce the the number of design iterations currently required down to a handful of forward passes (Norn et al., 2021). This would potentially come at the cost of using TPUs/GPUs with a larger memory than those used here.

Beyond improvements on the current method there are a myriad of ways that the current method could be expanded to tackle other design challenges and approaches. For example, functional design could be performed using infilling (Tischer et al., 2020) around some active site, where the sequence around some pre-specified active site is designed. The reverse could also be explored where the goal is to design some interface between two domains while leaving the backbone. The varying loss and confidence values that AlphaFold produces also provide additional ways that designs can be assessed. This could also be done in conjunction with statistical potentials (Tian et al., 2018) or molecular dynamics (Eastman et al., 2017). More broadly, models like AlphaFold open many new avenues for protein design methods, such as the work explored here, and will hopefully assist in paving the way for a new swathe of proteins with novel structures and functions.

## Acknowledgements

We thank members of the group for valuable discussions and comments.

